# The human cytomegalovirus-encoded pUS28 antagonizes CD4+ T-cell recognition by targeting CIITA

**DOI:** 10.1101/2023.10.17.562683

**Authors:** Fabienne Maaßen, Vu Thuy Khanh Le-Trilling, Luisa Betke, Thilo Bracht, Corinna Schuler, Malte Bayer, Antonia Belter, Tanja Becker, Benjamin Katschinski, Lori Frappier, Barbara Sitek, Katharina Fleischhauer, Mirko Trilling

## Abstract

Human cytomegalovirus (HCMV) is a relevant pathogen especially for individuals with impaired immunity. Harnessing potent immune antagonists, HCMV circumvents sterile immunity. Given that HCMV prevents the upregulation of *human leukocyte antigen* (HLA)-DP and HLA-DR, we screened a library of HCMV genes by co-expression with the HLA class II (HLA-II)-inducing transcription coordinator *class II transactivator* (CIITA). We identified the latency regulator pUS28 as interaction factor and potent viral antagonist of CIITA-driven expression of CD74, HLA-DR, HLA-DM, HLA-DQ, and HLA-DP. Both wt-pUS28 and a mutant incapable to induce G-protein-coupled signaling (R129A), but not a mutant lacking the C-terminus, drastically reduced the CIITA protein abundance post-transcriptionally. While control CD4+ T cells from HCMV-seropositive individuals vigorously responded to CIITA-expressing cells decorated with HCMV antigens, pUS28 expression was sufficient to inhibit HLA-II induction and immune recognition by HCMV-specific CD4+ T cells. Our data uncover a mechanism employed by HCMV to evade HLA-II-mediated recognition by CD4+ T cells.

## Introduction

More than half of the entire adult human population and over 90% of the elderly in developing countries are latently infected with the human cytomegalovirus (HCMV, human betaherpesvirus 5 [HHV-5]; NCBI: txid10359) ^1^. In healthy adults, HCMV raises strong canonical as well as non-canonical immune responses that limit viral replication and, in the vast majority of cases, prevent overt clinical manifestations, while leaving a substantial footprint on the immune system. Accordingly, more than half of all immune parameters are altered in HCMV-infected individuals^2^. Equipped with an astonishing arsenal of highly potent immune evasins, HCMV circumvents sterile immunity and instead establishes a life-long latency from which it reactivates during episodes of stress or impaired immunity. In cases of immune immaturity, immune senescence or impairment of the immune system, HCMV replication is not properly controlled, frequently resulting in severe morbidity and mortality in patients such as congenitally infected infants, people living with HIV, and transplant recipients ^3^.

Although the concerted activity of various mediators of innate and adaptive immunity is crucial for efficient cytomegalovirus (CMV) control ^4^, two aspects are particularly important: interferon (IFN) responses and CD4+ T cells. Accordingly, the depletion of CD4+ cells together with IFNγ results in CMV reactivation in more than 75% of animals ^4^. IFN secretion is among the first responses elicited upon pathogen encounter. IFNs trigger signaling cascades and initiate a specific transcriptional profile that fosters intrinsic immunity, induces direct innate immunity, and orchestrates adaptive immune responses. In this regard, IFNs act by inducing IFN-stimulated genes (ISGs), many of which being known for their antiviral activity, and by down-regulating IFN-repressed genes (IRepGs) ^5–7^. In addition to their ability to establish a cell-intrinsic antiviral state, IFNs booster adaptive immune responses. In particular, IFNγ is well known as key cytokine for antiviral Th1 responses and as strong enhancer of antigen presentation. In accordance with the relevance of IFNs, the absence of IFN-induced signaling renders mice extremely vulnerable to CMV infections ^8–10^. CMV infection induces strong CD4+ T-cell responses. While best known for their cytokine secretion helping B cells and CD8+ T cells, CD4+ T cells can elicit antiviral activity ^11–13^. The latter is associated with the ability to secrete antiviral cytokines such as IFNγ. Furthermore, certain CD4+ T cells can kill virus-infected cells ^14^. HCMV-specific CD4+ T cells were shown to produce IFNγ, TNFα, and granzyme B ^15^ and to lyse HCMV antigen-expressing cells *in vitro* ^16,17^. Accordingly, CD4+ T-cell-mediated immunity is crucial for CMV control in mouse models *in vivo* ^11,18,19^, and for controlling maternal viremia and prevention of severe CMV-associated fetal disease during primary rhesus CMV infections ^20^. Furthermore, CD4+ T cells are fundamental to prevent HCMV reactivation in immunocompromised patients receiving solid organ allografts ^12^ or hematopoietic cell transplantation (HCT), while insufficient CD4+ T-cell levels in transplant recipients are associated with recurrent HCMV reactivation, end-organ disease, and an increased likelihood of lethal infections ^21,22^.

For the canonical T-cell receptor (TCR)-dependent activation of CD4+ T cells, antigenic peptides must be presented by the HLA-II molecules HLA-DR, HLA-DQ or HLA-DP. While antigen-presenting cells (APCs) constitutively express HLA-II, various other cell types start to express HLA-II when they are exposed to IFNγ ^23–26^. Activated CD4+ T cells efficiently produce IFNγ, leading to a positive feed-forward loop of increased HLA-II presentation and enhanced CD4+ T-cell recognition in the infected niche. Constitutive as well as IFNγ-induced HLA-II expression are both mediated by the class II transactivator (CIITA), which is the essential and sufficient master regulator of the transcription of the genes in the HLA-II locus. Accordingly, CIITA and its co-factors control the expression of the classical HLA-II molecules DR, DQ, DP, the non-classical HLA-II peptidome editors DM and DO, and the invariant chain (Ii, also known as CD74) ^26,27^. The absence of functional CIITA results in a loss of HLA-II presentation, leading to a hereditary immunodeficiency called type II bare lymphocyte syndrome (BLS) that causes an extreme vulnerability to infections. In accordance with the relevance of CD4+ T cells and CIITA-driven HLA-II presentation, BLS patients frequently suffer from severe, persistent HCMV infections ^28,29^.

For allogenic hematopoietic cell therapy (HCT), the HLA-DP locus is of particular interest given the frequent incompatibility of patients and donors ^30^, the allotype dependency of peptide presentation and recognition by alloreactive CD4+ T cells ^31–33^, and the relevance for chronic virus infections as indicated by the strong association of HLA-DPB1 SNPs with chronic hepatitis B virus (HBV) infections (see e.g., ^34^). CD4+ T cells recognizing HLA-DP-restricted peptides have also been described for HCMV ^35–39^, but to our knowledge, viral countermeasures have not yet been described.

HCMV-propagating cells such as human fibroblasts, endothelial cells, and epithelial cells are capable to induce HLA-II presentation after exposure to IFNγ. Moreover, cells serving as a reservoir for HCMV latency, e.g., monocytes, constitutively express HLA-II. CD4+ T cells can recognize HCMV-derived antigens known to be expressed during latency, leading to the recognition of latently infected cells and the production of IFNγ and/or cytotoxic responses ^40^. These facts raise the question, how HCMV circumvents CD4+ T-cell-mediated elimination during productive replication as well as latency.

Here, we show that the HCMV-encoded G-protein-coupled receptor (GPCR) pUS28 directly targets CIITA, impairing HLA-II presentation and CD4+ T-cell recognition. Importantly, pUS28 is among the few viral proteins abundantly expressed during both productive replication as well as experimental and natural latency ^41–45^. Thus, our data reveal a novel mechanism employed by HCMV to circumvent the recognition by CD4+ T cells during different stages of infection.

## Results

### HCMV counteracts IFN_γ_-induced HLA-DP induction

Given the aforementioned relevance of HLA-DP in HCT and chronic virus infections, we interrogated if and how HCMV affects HLA-DP induction and presentation. IFNγ induced the upregulation of HLA-DP on the surface of HCMV-permissive fibroblasts (Fig. 1a and 1b), whereas this was not the case after treatment with IFNα or TNFα (Fig. 1b). Longer periods of IFNγ incubation led to an increase of HLA-DP and HLA-DR on the cell surface (Fig. 1c). In clear contrast to uninfected control cells, the upregulation of HLA-DP by IFNγ was prevented in HCMV-infected cells (Fig. 1d and 1e). Interestingly, the same inhibitory phenotype was observed upon infection with HCMV mutants lacking the *US2-6* or the *US2-11* gene region (Fig. 1d and 1e), that comprise the genes for pUS2 and pUS3, which degrade HLA-DRα and HLA-DMα ^46^ and block the assembly of HLA-DRα/β heterodimers ^47^, respectively. These data indicate that HCMV encodes so far unknown antagonists of IFNγ-induced HLA-DP presentation located outside of the *US2-11* gene region.

**Figure 1:**
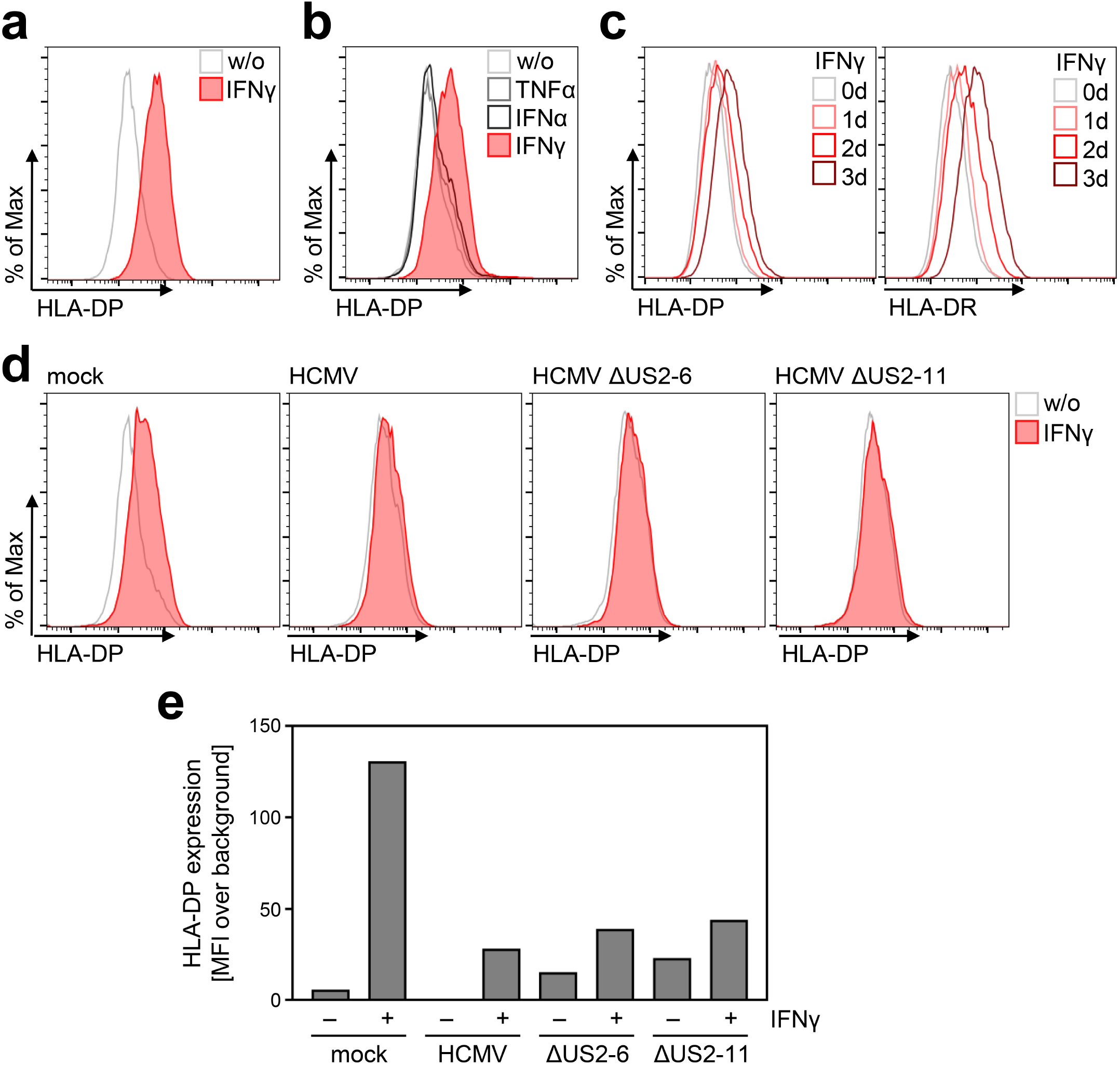
IFN_γ_-induced HLA-DP expression is abrogated in HCMV-infected fibroblasts. (a) MRC-5 fibroblasts either were left untreated or were treated with 200 U/ml IFNγ. At 72 h post-treatment, cells were stained with anti-HLA-DP antibody and analyzed by flow cytometry. w/o, untreated. (b) MRC-5 fibroblasts either were left untreated or were treated with 200 U/ml IFNγ, 200 U/ml IFNα, or 20 ng/ml TNFα. At 48 h post-treatment, cells were stained with anti-HLA-DP antibody and analyzed by flow cytometry. w/o, untreated. (c) MRC-5 fibroblasts either were left untreated or were treated with 200 U/ml IFNγ for 1, 2, or 3 days. Cells were stained with anti-HLA-DP or anti-HLA-DR antibody and analyzed by flow cytometry. (d) MRC-5 fibroblasts were either mock infected or were infected (MOI 3) with AD169 (HCMV), AD169-BAC2 (HCMVΔUS2-6), or AD196-BAC2ΔUS2-11 (HCMVΔUS2-11). At 4 h post-infection, cells were treated with 200 U/ml IFNγ. After 48 h of treatment, cells were stained with anti-HLA-DP antibody and analyzed by flow cytometry. w/o, untreated. (e) The mean fluorescence intensity (MFI) values of HLA-DP expression of untreated or IFNγ-treated MRC-5 cells, mock treated or infected (as in Fig. 1d).

### The HCMV-encoded pUS28 antagonizes CIITA-induced HLA-II expression

Since HCMV prevented HLA-DP induction, we aimed at identifying the responsible gene product(s). To this end, we applied an expression library, which has been previously described ^48^, comprising more than 150 canonical HCMV genes to screen for HLA-DP antagonists. To minimize potentially confounding effects of viral antagonists of IFN signaling acting upstream of CIITA induction, we set up the screen by co-transfecting a CIITA expression plasmid together with individual vectors encoding HCMV proteins. Afterwards, the CIITA-induced HLA-DP cell surface expression was quantified by cytometry (Fig. 2a). The screen revealed candidates for the observed antagonism of HLA-DP induction including pUS9, pUS19, pp65-UL83, and pUS28. Validation experiments substantiated that pUS28 was reproducibly capable to significantly diminish CIITA-induced HLA-DP surface expression (Fig. 2b and data not shown). Due to its relevance for the HCMV biology, we further assessed the influence of pUS28 on the CIITA-dependent induction of other genes in the HLA-II locus. Similar to its effect on HLA-DP, pUS28 also significantly diminished the cell surface expression of the presenting molecules HLA-DR (Fig. 2c and 2d) and HLA-DQ (Fig. 2d) as well as the peptide editor HLA-DM (Fig. 2d). Given that different cell types express varying levels of CIITA, we tested the dose-response relationship between CIITA and pUS28. While increasing amounts of CIITA dose-dependently drove HLA-DP and HLA-DR expression in transfected cells, pUS28 significantly and dose-dependently inhibited surface levels of HLA-DR (Fig. 2e) and HLA-DP (Fig. 2f), irrespective of the CIITA abundance.

**Figure 2:**
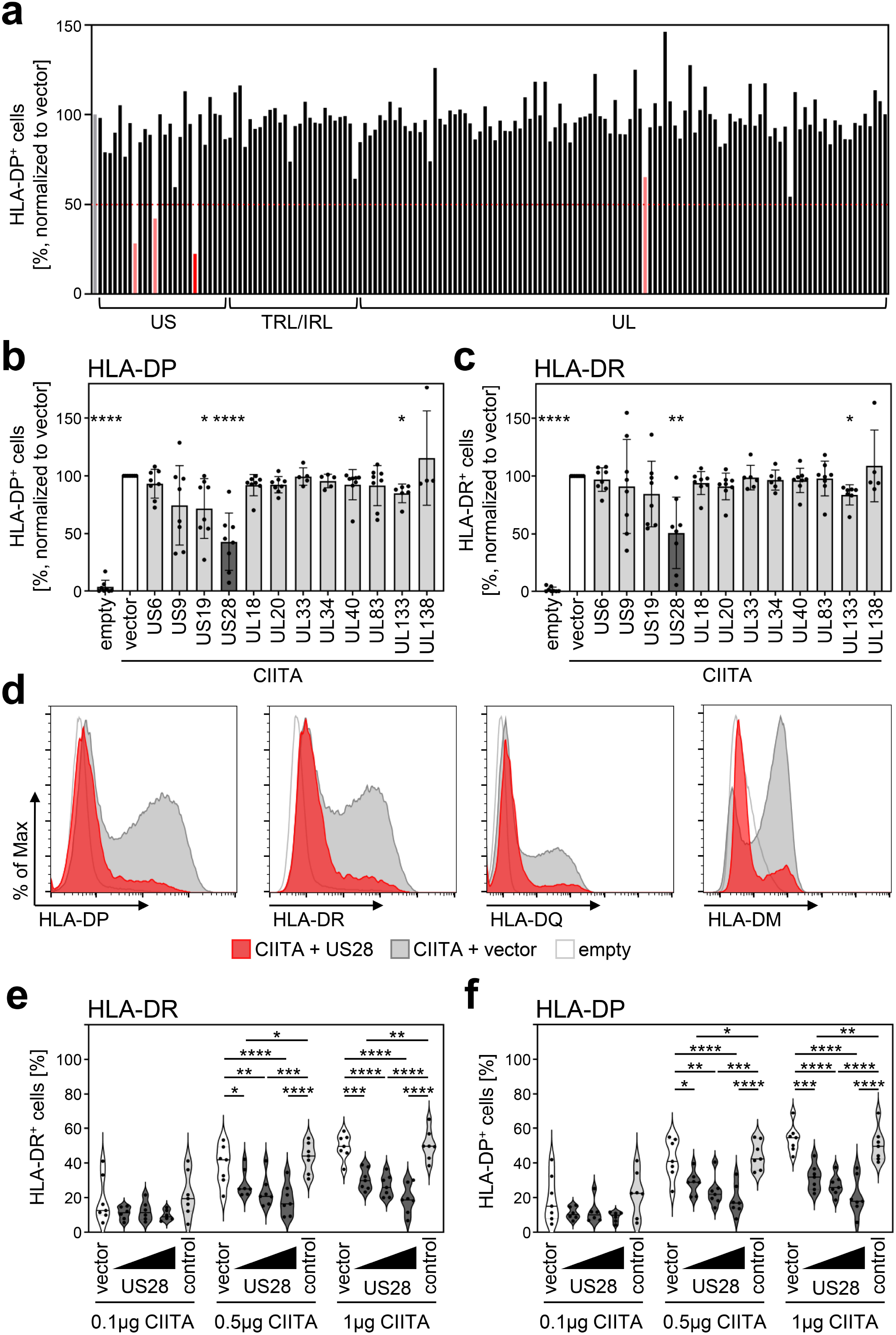
HCMV-pUS28 functions as antagonist of the CIITA-induced HLA class II upregulation. (a) HeLa cells were co-transfected with a CIITA expression construct and a library of single HCMV-gene encoding plasmids or empty vector. At 48 h post-transfection, cells were stained with anti-HLA-DP antibody and analyzed by flow cytometry. Cell surface expression of HLA-DP was normalized to cells transfected with CIITA expression construct and empty vector. The percentage of HLA-DP-positive cells is shown. (b/c) HeLa cells were co-transfected with a CIITA expression construct and indicated plasmids or empty vector. At 48 h post-transfection, cells were stained with anti-HLA-DP (b) or anti-HLA-DR (c) antibody and analyzed by flow cytometry. Cell surface expression of HLA class II was normalized to cells transfected with CIITA expression construct and empty vector. The percentage of HLA-DP- or HLA-DR-positive cells is shown (mean values ± SD, n = 4-8). Significance was calculated by Kruskal-Wallis test compared to empty vector control. Empty, untransfected. Vector, empty vector control. (d) HeLa cells were either left untreated or were co-transfected with a CIITA expression construct and pcDNA:US28-HA or empty vector. At 48 h post-transfection, cells were stained with anti-HLA-DP, anti-HLA-DR or anti-HLA-DQ antibody or intracytoplasmic stained with anti-HLA-DM antibody and analyzed by flow cytometry. (e/f) HeLa cells were co-transfected with indicated amounts of CIITA expression construct and increasing doses of pcDNA:US28-HA, a control plasmid (pcDNA:US29-HA) or empty vector. The total DNA amount of each transfection was adjusted to the same level by adding the respective amount of empty vector. At 48 h post-transfection, cells were stained with anti-HLA-DR (e) and anti-HLA-DP (f) antibodies and analyzed by flow cytometry. The percentage of HLA-DR- and HLA-DP-positive cells is shown (n = 5-8). Significance was calculated by two-way ANOVA test. Vector, empty vector control. Control, control plasmid.

### pUS28 down-regulates CIITA post-transcriptionally

To probe into the underlying molecular mechanism of the pUS28-mediated HLA-II inhibition, untagged or epitope-tagged CIITA were co-expressed with pUS28 or a control protein. Afterwards, transcript levels of *CIITA* were quantified by semi-quantitative reverse transcriptase (RT)-PCR. Irrespective of the presence or absence of pUS28, the levels of tagged as well as untagged *CIITA* mRNA remained unaltered (Fig. 3a), while the amounts of HLA-DP and HLA-DR mRNA were decreased (data not shown). In contrast to the unchanged mRNA levels of *CIITA*, a parallel evaluation of CIITA protein amounts revealed a drastic reduction upon pUS28 co-expression (Fig. 3b), suggesting a post-transcriptional effect of pUS28 on CIITA. The CIITA down-modulating capacity of pUS28 was observed with different plasmid preparations (Fig. 3c), and was specific for CIITA, since control proteins such as the enhanced yellow fluorescent protein (EYFP) remained unaffected, arguing against a general influence of pUS28 on transcription or translation (Fig. 3c). The effect on CIITA was also not an overarching capacity of viral chemokine receptor homologues, since the protein pUS27, a GPCR encoded by the neighboring gene in the viral genome, did not decrease CIITA protein amounts (Fig. 3d) or HLA-DP surface levels (Fig. 3e).

**Figure 3:**
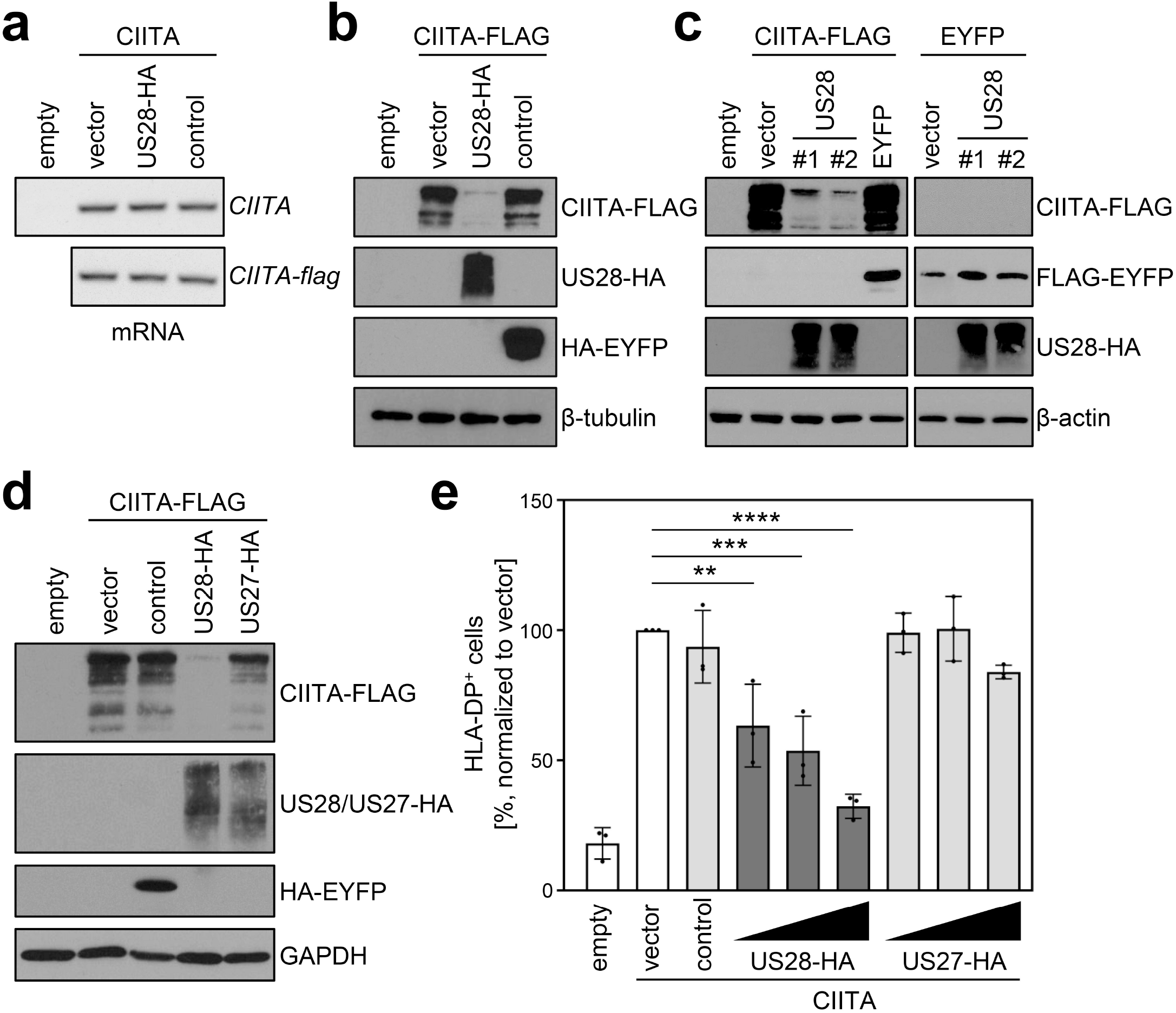
pUS28 down-regulates CIITA post-transcriptionally. (a/b) HeLa cells were either left untreated or were co-transfected with CIITA or CIITA-3xFLAG expression constructs and pcDNA:US28-HA, pIRESNeo-FLAG/HA-EYFP (control) or empty vector. At 24 h post-transfection, cells were harvested and split for preparation of total RNA and protein lysate. RNA samples were used for semi-quantitative RT-PCR with indicated gene-specific primers (a) and protein lysates were analyzed by immunoblot using antibodies detecting the indicated proteins or the respective epitope tags (b). (c) HeLa cells either were left untreated or were co-transfected with the indicated plasmids. At 24 h post-transfection, protein lysates were generated and analyzed by immunoblot using antibodies detecting the indicated proteins or the respective epitope tags. #1, #2, different plasmid preparations. (d) HeLa cells were either left untreated or were co-transfected with CIITA-3xFLAG expression construct and pcDNA:US28-HA, pcDNA:US27-HA, pIRESNeo-FLAG/HA-EYFP (control) or empty vector. At 24 h post-transfection, protein lysates were generated and analyzed by immunoblot using antibodies detecting the indicated proteins or the respective epitope tags. (e) HeLa cells were co-transfected with CIITA expression construct and increasing doses of pcDNA:US28-HA or pcDNA:US27-HA, pIRESNeo-FLAG/HA-EYFP (control) or empty vector. The total DNA amount of each transfection was adjusted to the same level by adding the respective amount of empty vector. At 48 h post-transfection, cells were stained with anti-HLA-DP antibody and analyzed by flow cytometry. The percentage of HLA-DP-positive cells normalized to empty vector control is shown. Empty, untransfected. Vector, empty vector control. Control, pIRESNeo-FLAG/HA-EYFP.

### Global mass spectrometry confirmed the pUS28-mediated decrease of the CIITA abundance and identified downstream targets of the HLA class II pathway

To address the influence of pUS28 on CIITA and the proteome, we performed global mass spectrometry (MS) analyses in which we compared control cells with cells expressing either pUS28, CIITA or both (Fig. 4a and 4b, Supplementary Table S1). As expected, our MS analyses confirmed the expression of pUS28 (Fig. 4a-c). In agreement with previous data, pUS28 led to a significant upregulation of IL-6 ^49^, and attenuated AP-1 components^50^ (here JunD) (Fig. 4d and 4e), validating our experimental setup. These unbiased MS analyses confirmed the significant downregulation of CIITA by pUS28 (Fig. 4f). Other components of the CIITA enhanceosome such as RFX5, RFX-AP, NF-YA, and NF-YC (Supplementary Fig. S1), as well as recently described factors^51^ that influence HLA-II transcription showed constitutive, CIITA-independent expression (Supplementary Fig. S2). Conversely, CD74 - the invariant chain required for proper maturation and loading of HLA class II molecules - was significantly induced by CIITA, unless pUS28 was co-expressed (Fig. 4g). The negative effect of pUS28 on CIITA-driven CD74 expression was confirmed by flow cytometry (Fig. 4h). In contrast to several other CIITA-regulated genes, CD74 is not encoded by the HLA-II locus on chromosome 6. Thus, the fact that pUS28 prevents CIITA-induced CD74 induction suggests that pUS28 elicits the negative effect on CIITA target genes independent of their genomic localization in the MHC-II locus.

**Figure 4:**
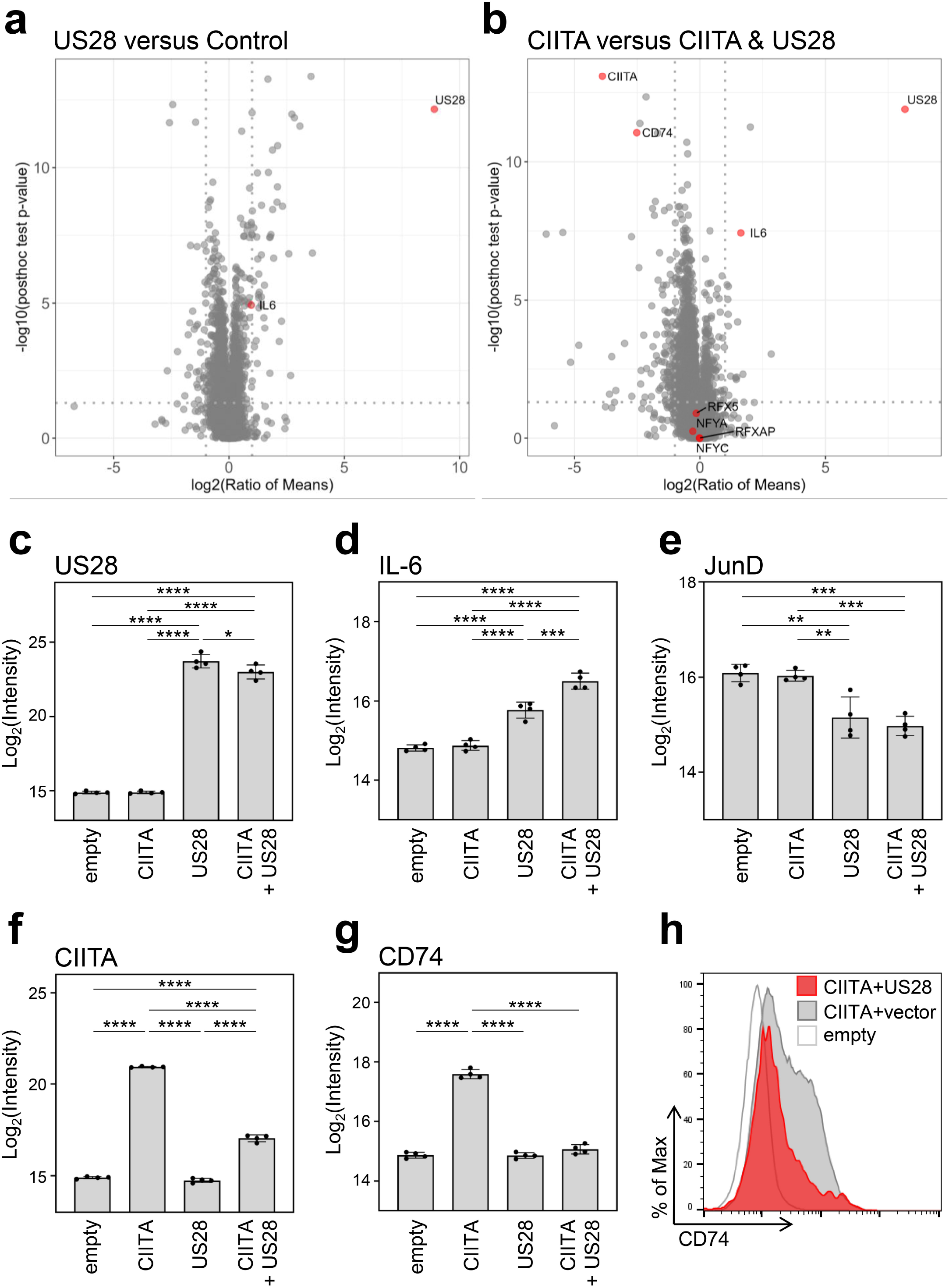
Global proteome analysis revealed that pUS28 targets CIITA and affects the CIITA-regulated protein CD7. (a/b) HeLa cells were either left untreated or were transfected with CIITA-3xFLAG expression construct, pcDNA:US28-HA or both. At 24 h post-transfection, whole cell lysates were generated and subjected to mass spectrometric analysis of protein abundance. Volcano plots showing log2 (ratio of means) (x-axis) versus significance (y-axis) of the comparison of untreated and pUS28-expressing cells (a) or cells expressing CIITA in the presence or absence of pUS28 (b). Proteins are indicated as grey dots, highlighted in red are IL-6 (known to be upregulated by pUS28), co-factors of HLA-II transcription and signaling, CIITA, and pUS28. (c-g) Changes in the abundance of selected proteins detected by MS: (c) pUS28, (d) IL-6, (e) JunD, (f) CIITA, (g) CD74. Depicted are log2 (intensity) values of untreated cells, cells expressing either CIITA, pUS28 or both (n = 4). (h) HeLa cells were either left untreated or were co-transfected with a CIITA expression construct and pcDNA:US28-HA or empty vector. At 48 h post-transfection, cells were stained with anti-CD74 antibody and analyzed by flow cytometry. Empty, untransfected. Vector, empty vector control.

### pUS28 interacts with CIITA and reduces its half-life irrespective of the G protein-coupling capacity

Next, we tested if pUS28 and CIITA physically interact. To circumvent the issue that CIITA levels were diminished to almost undetectable levels upon pUS28 co-expression, lysates either containing pUS28 or CIITA were combined before an immunoprecipitation (IP) was performed. The IP of pUS28 co-purified CIITA, indicating that both proteins form physical complexes (Fig. 5a). To test if pUS28 forces CIITA into a detergent-insoluble fraction, cell lysates were prepared with a denaturing lysis buffer (based on high urea concentrations) and subjected to immunoblot analysis. This approach did not lead to the reappearance of CIITA (Supplementary Fig. S3). Irrespective of the absence or presence of pUS28, we could not detect CIITA in cell culture supernatants (data not shown), arguing against pUS28-mediated CIITA shedding. Since we did not find evidence for sequestration or shedding, we tested for a pUS28-mediated CIITA degradation. To this end, we compared the half-life of CIITA in the presence or absence of pUS28. Despite the inherently short half-life of CIITA ^52^, we observed a more rapid CIITA decay when pUS28 was co-expressed (Fig. 5b).

**Figure 5:**
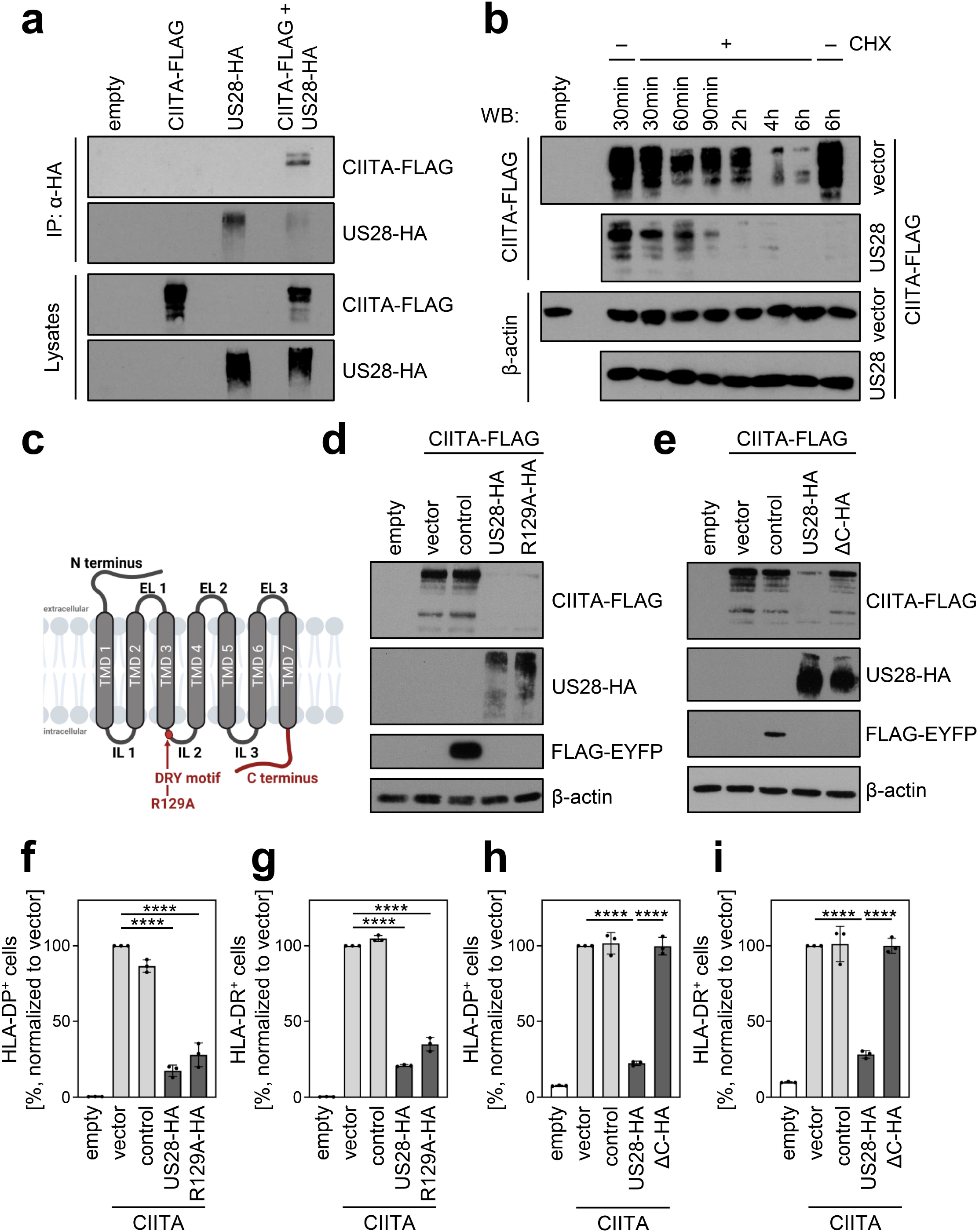
pUS28 reduces the half-life of CIITA, independently of the G protein-coupling capacity. (a) HeLa cells were either left untreated or were transfected with CIITA-3xFLAG expression construct or pcDNA:US28-HA. At 24 h post-transfection, protein lysates were generated and an IP with an HA-specific mouse monoclonal antibody was performed either with unmodified samples or with mixed samples of CIITA- and US28-transfected cells. The lysates and IP samples were analyzed by immunoblot to test CIITA co-precipitation. (b) HeLa cells were co-transfected with CIITA-3xFLAG expression construct and pcDNA:US28-HA or empty vector. At 16 h post-transfection, cells either were left untreated or were incubated with 50 µg/ml cycloheximide (CHX) for indicated periods. Protein lysates were generated and analyzed by immunoblot using antibodies detecting the indicated proteins or the respective epitope tags. (c) Schematic overview of the structure of pUS28. Indicated are all structural parts of the protein and the DRY motif. Mutation of the arginine in this motif to alanine (R129A) ablates G-protein coupling. Created with BioRender.com. (d) HeLa cells were either left untreated or were co-transfected with CIITA-3xFLAG expression construct and pcDNA:US28-HA, pcDNA:US28-R129A-HA, pIRESNeo-FLAG/HA-EYFP or empty vector. At 24 h post-transfection, protein lysates were generated and analyzed by immunoblot using antibodies detecting the indicated proteins or the respective epitope tags. (e) HeLa cells were either left untreated or were co-transfected with CIITA-3xFLAG expression construct and pcDNA:US28-HA, pcDNA:US28-ΔC-HA, pIRESNeo-FLAG/HA-EYFP (control) or empty vector. At 24 h post-transfection, protein lysates were generated and analyzed by immunoblot using antibodies detecting the indicated proteins or the respective epitope tags. (f/g) HeLa cells were either left untreated or were co-transfected with CIITA expression construct and pcDNA:US28-HA, pcDNA:US28-R129A-HA, pIRESNeo-FLAG/HA-EYFP (control) or empty vector. At 48 h post-transfection, cells were stained with anti-HLA-DP (f) or anti-HLA-DR (g) antibodies and analyzed by flow cytometry. Cell surface expression of HLA-DP or HLA-DRwas normalized to cells transfected with CIITA expression construct and empty vector. The percentage of HLA-DP- and HLA-DR-positive cells is shown (n = 3). (h/i) HeLa cells were either left untreated or were co-transfected with CIITA-3xFLAG expression construct and pcDNA:US28-HA, pcDNA:US28-ΔC-HA, pIRESNeo-FLAG/HA-EYFP (control) or empty vector. At 48 h post-transfection, cells were stained with anti-HLA-DP (h) or anti-HLA-DR (i) antibodies and analyzed by flow cytometry. Cell surface expression of HLA-DP or HLA-DR was normalized to cells transfected with CIITA expression construct and empty vector. The percentage of HLA-DP- and HLA-DR-positive cells is shown (n = 3). Empty, untransfected. Vector, empty vector control. ΔC, C-terminal (aa 298-354) deletion mutant of pUS28. R129A, point mutation mutant of pUS28. Control, pIRESNeo-FLAG/HA-EYFP.

Intriguingly, a panel of inhibitors targeting different cellular degradation pathways (e.g., the neddylation inhibitor MLN4924, the inhibitor of the proteasome MG-132, the autophagy inhibitors bafilomycin and 3-MA, the inhibitors of lysosomal acidification chloroquine and ammonium chloride, the protease inhibitors E-64 and pepstatin A, the caspase inhibitor zVAD-FMK as well as the convertase inhibitor Decanoyl-RVKR-CMV) all failed to restore CIITA levels in pUS28-expressing cells (data not shown), implying that pUS28 acts either by redundant or unusual degradation processes.

To assess if the G-protein signaling of pUS28 is essential for the down-regulation of CIITA, we compared wt-pUS28 and R129A-pUS28. The latter is a well-studied mutant of pUS28 that harbors a single amino acid substitution at position 129 (arginine to alanine) within the canonical DRY motif, abrogating G-protein signaling without compromising the subcellular localization or internalization (^53^; visualized in Fig. 5c). Both pUS28 variants diminished CIITA protein levels (Fig. 5d). Conversely, the intracellular C terminus turned out to be indispensable for the CIITA degradation (Fig. 5e). In accordance with these findings, R129A-pUS28 and wt-pUS28 similarly diminished the surface expression of HLA-DP (Fig. 5f) and HLA-DR (Fig. 5g), while the mutant lacking the C terminus did not (Fig. 5h and 5i), indicating that pUS28 targets CIITA through its C terminus.

### pUS28 antagonizes HCMV-specific CD4+ T cells

After identification of pUS28 as viral antagonist of the CIITA-driven HLA class II expression and having in mind that pUS28 is expressed during HCMV latency, we aimed to investigate the relevance of pUS28 for immune recognition. HCMV-specific CD4+ T cells were enriched by antigen-dependent expansion using peripheral blood mononuclear cells (PBMCs) from HCMV-seropositive healthy individuals exposed to lysates derived from HCMV-infected fibroblasts (see schematic overview in Fig. 6a). During the 24 h re-stimulation phase, the HCMV-specific CD4+ T cells were co-cultured with HCMV antigen-loaded fibroblasts expressing CIITA together with pUS28, with pUS28-R129A or with an irrelevant protein. CD4+ T-cell activation was evaluated by CD137 upregulation. HCMV-specific CD4+ T cells from HCMV-positive individuals vigorously responded to CIITA-expressing cells, but not to control cells that did not express CIITA (Fig. 6b), indicating a specific HLA-II-dependent T-cell activation. In the presence of pUS28 or the mutant R129A-pUS28, the immune recognition of CIITA-expressing, antigen-loaded cells by HCMV-specific CD4+ T cells was significantly reduced (Fig. 6b). Since CD4+ T cells are well known for the ability to produce antiviral cytokines such as IFNγ, we assessed the cell culture supernatants collected after the 24 h re-stimulation phase regarding their antiviral activity against HCMV. We conditioned human fibroblasts with the supernatants prior to the infection with an EGFP-expressing reporter HCMV. Afterwards, we quantified HCMV-induced EGFP expression (Fig. 6c) and visualized the degree of HCMV infection by microscopy (Fig. 6d). Supernatants from HCMV-specific CD4+ T cells stimulated with CIITA-expressing cells strongly inhibited HCMV-induced EGFP expression, while supernatants derived from CD4+ T cells stimulated in the presence of pUS28 showed diminished antiviral activity (Fig. 6c and 6d). These data demonstrate that pUS28 inhibits the CIITA-driven and HLA-II-dependent CD4+ T-cell activation in terms of the production of antiviral cytokines.

**Figure 6:**
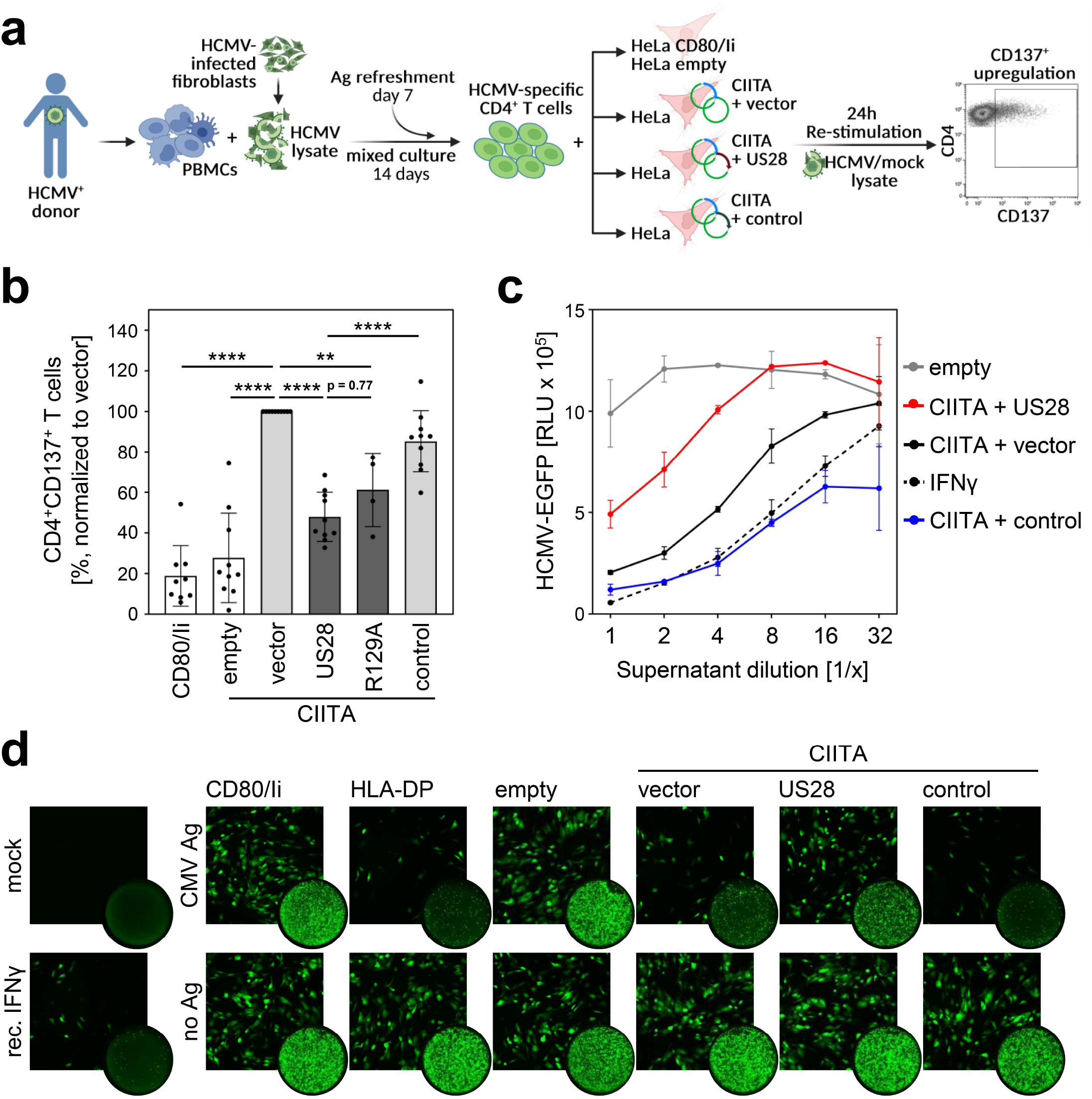
Activation of HCMV-specific CD4+ T cells is inhibited by pUS28. (a) Schematic overview of the experimental setup. PBMCs of HCMV-seropositive healthy donors were isolated, pulsed with HCMV lysate and incubated for 14 days, with an antigen refreshment step at day 7. Afterwards, cells were co-cultured with HeLa cells that were either left untreated or were co-transfected with CIITA expression construct and pcDNA:US28-HA, pcDNA:US28-R129A-HA, pIRESNeo-FLAG/HA-EYFP or empty vector, 48 h prior to co-culture, and were re-stimulated with mock or HCMV lysate. HeLa cells only expressing CD80 and the invariant chain (HeLa CD80/Ii) served as further negative control. After 24 h of incubation, the specific T-cell response was quantified by flow cytometry as percentage of gated CD4+ T cells expressing the activation marker CD137 (bottom). Created with BioRender.com. (b) Activation of HCMV-specific CD4+ T cells was measured as described in (A). Proportion of CD137-positive T cells was normalized to T cells activated by HeLa cells transfected with CIITA expression construct and empty vector, and pulsed with HCMV lysate. Mean values ± SD are depicted (n = 4-10). Significance was calculated by one-way ANOVA test. (c) MRC-5 cells were incubated with supernatants from HCMV-specific CD4+ T cells (B) or recombinant IFNγ in serial dilutions for 24 h. Next, cells were infected with BAC20-EGFP at an MOI of 0.05 and HCMV-induced EGFP expression was measured at 5 days post-infection. (d) MRC-5 cells were incubated and infected as in (c) and infected cells were visualized by fluorescence microscopy after 4 days of infection (square pictures) or whole-well imaging (circle pictures) at 6 days post-infection. Empty, untransfected. Vector, empty vector control. R129A, point mutation mutant of pUS28. Control, pIRESNeo-FLAG/HA-EYFP. Mock, uninfected. CMV Ag, HCMV lysate-treated. No Ag, mock lysate-treated.

## Discussion

In this study, we identified pUS28 as HCMV-encoded antagonist of CIITA and CIITA-driven HLA class II expression (Fig. 7). The viral protein was found to physically interact with CIITA, causing a post-transcriptional decline of the CIITA protein by an increased protein decay. This CIITA degradation was sufficient to decrease HLA-DR, HLA-DQ, HLA-DM, CD74/invariant chain, and HLA-DP cell surface expression and to abrogate activation of HCMV-specific CD4+ T cells. Despite its role as a highly relevant human pathogen, attenuated HCMVs became the basis for the development of promising vaccine vectors (e.g., for the vaccination against lentiviruses ^54^), which may benefit from the deletion of immune evasins such as US28, either in terms of enhanced immunogenicity and/or increased safety.

Briefly after its discovery as HCMV-encoded GPCR, its functionality as calcium-mobilizing beta chemokine receptor was documented ^55^. Since then, pUS28 became one of the best-studied HCMV proteins, and several important immunological and viral functions have been assigned to this molecule ^44^. In terms of its expression profile, pUS28 is special due to its presence in experimental and natural latency ^41–45^, constituting a drug target for the latent HCMV reservoir using specifically designed toxins ^56^. Our study adds the inhibition of CIITA-induced HLA-II antigen presentation to the list of important pUS28 functions. The relevance of pUS28 for latency establishment and reactivation has been studied in huNSG mice ^57^. These mice develop certain aspects of a human T-cell compartment ^58^, which may lead to HLA-II restricted elimination of infected cells by CD4+ T cells. In addition to its influence on latency and reactivation, the herein documented ability of pUS28 to counteract CIITA-driven HLA-II presentation may influence the outcome of such experiments by effects of pUS28 on the CD4+ T-cell recognition by the immune system. Similarly, pUS28-expressing mouse cytomegalovirus (MCMV) mutants have been studied in mice ^59^. In this regard, it is important to note that pUS28 is capable to down-regulate mouse CIITA (Supplementary Fig. S4).

Elegant work by the Sinclair laboratory revealed that pUS28, besides its function as GPCR and latency regulator, targets certain host proteins such as MNDA/PYHIN3 and IFI16/PYHIN2 for rapid degradation. In contrast to the effect on CIITA shown here, however, this degradation was dependent on GPCR signaling as indicated by the restoration of target protein levels when R129 was mutated ^60^. Although CIITA and HLA-DP levels were neither assessed nor discussed, the same work documented a negative effect of pUS28 on constitutive HLA-DR expression, while no difference between wt-pUS28 and an R129A mutant with regard to the IFNγ-induced HLA-DR expression was observed ^60^. Importantly, Elder *et al*. and Slobedman *et al.* showed that latently infected primary human CD14+ monocytes and human fetal liver hematopoietic cells exhibit less HLA-DR on the surface compared to uninfected cells ^60,61^. Our data regarding the targeting of CIITA by pUS28 now provide a parsimonious molecular explanation for these findings, and extend it to other highly relevant HLA-II molecules including CD74, HLA-DM, HLA-DQ, and HLA-DR.

Recent work with the MCMV model showed that the deletion of the *US28* homologue *M33* results in altered MHC-I presentation in an allotype-specific manner (H-2L^d^ and H-2K^d^ being down-regulated in an *M33*-dependent manner, but not H-2D^d^) ^62^. Our data raise the intriguing question, if pM33, like pUS28, affects CIITA and the downstream expression of murine MHC-II molecules.

Previous studies focused on the interplay between HCMV infection and HLA-DR presentation, largely neglecting other HLA-II molecules such as HLA-DP. Work by Miller *et al*. showed that HCMV counteracts IFNγ-induced gene expression, partly by inducing the proteasomal degradation of JAK1, leading to diminished CIITA and HLA-DRα induction when HCMV-infected cells are exposed to IFNγ ^63^. Furthermore, HCMV targets constitutive as well as induced CIITA-dependent HLA-II expression at multiple levels by a multipronged attack, comprising the inhibition of (I) IFN-JAK-STAT signaling ^63–65^, (II) constitutive CIITA transcription ^66^, (III) CIITA protein stability (as shown here), and (IV) HLA-DR degradation and translocation ^46,47,67^. This high level of redundancy may also explain why others concluded, based on loss-of-function experiments, that the *US* gene region comprising *US28* is dispensable for the inhibition of HLA-DR presentation ^68^, while we observed a clear gain-of-function regarding HLA-II inhibition upon pUS28 expression. Accordingly, a BACmid-derived ΔUS28-HCMV, which also lacks the *US2-US6* gene region due to the insertion of the BAC cassette, still inhibited HLA-II presentation (data not shown) indicating the existence of additional HLA-II antagonists.

HLA-II-restricted CD4+ T-cell immunity to HCMV is crucial for the control of the lifelong infection by this virus. Its impairment in immunocompromised patients, including HCT recipients, is associated with considerable clinical risks. The present study is the first to identify a mechanism by which HCMV down-regulates HLA-DP during latency, although it is likely that the abovementioned alternative pathways employed by HCMV to reduce HLA-II levels also affect HLA-DP.

In our introduction, we eluded to the wealth of knowledge regarding the critical importance of CD4+ T cells for the immune control of cytomegaloviruses in mouse and rhesus models as well as in humans. These findings seem to contradict the multitude of CMV-encoded inhibitors of constitutive and induced HLA-II presentation. How can these two, at first glance mutually exclusive, facts be reconciled? The first argument is that cytomegaloviruses and their hosts are situated in an evolutionary red-queen race that establishes a hard-fought equilibrium. Thus, the residual CD4+ T-cell-mediated immune control recognizing CMV-infected cells presenting diminished HLA-II levels may still be crucial for host survival despite the existence of viral inhibitors. Another intriguing possibility is that HLA-II-mediated CD4+ T-cell activation might be elicited by cells that do not become infected by HCMV such as NKG2C+ memory NK cells ^69^, which have been shown to be mediators of viral control in transplanted patients ^70^. The elucidation of the complex molecular mechanisms governing HCMV immune evasion in the immunocompetent and the immunocompromised host will provide important guidance for the design of tailored protocols of risk protection, e.g., by vaccination, targeted cellular therapies or drugs that interfere with pUS28-mediated CIITA degradation.

## Supporting information

Suppl.-Table S2 Primers

Suppl.-Table S1 Proteomics

## Acknowledgments

We thank Lejla Timmer, Kerstin von Ameln, and Sophie Eppler for excellent technical support, and the teams of the Fleischhauer and Trilling laboratories for insightful discussions. M.T. received funding from the Deutsche Forschungsgemeinschaft (DFG) through grants TR 1208/1-1 and TR 1208/2-1. KF received funding from the DFG through grant FL 843/1-1, the Deutsche José Carreras Leukämie Stiftung (DJCLS 20R/2019), the Dr. Werner Jackstädt Stiftung, and the Joseph Senker Stiftung.

## Author contributions

FM, VTKL-T, LB, CS, AB, BK, and TBe performed research. TBr, MB, and BS performed MS analysis. LF provided an indispensable resource (the expression library). FM, VTKL-T, KF, and MT interpreted data. VTKL-T, KF, and MT supervised the project. KF and MT raised funding for the project. FM, VTKL-T, and MT wrote the manuscript. All authors contributed to the article and approved the submitted version.

## Methods

### Cells and cell lines

HeLa cells (ATCC CCL-2), HeLa CD80/Ii ^71^, BJ-5ta cells (ATCC CRL-4001) and MRC-5 fibroblasts (ATCC CCL-171) were grown in Dulbecco modified Eagle medium (DMEM, Gibco) supplemented with 10 % (v/v) FCS (Sigma-Aldrich), 100 μg/ml streptomycin/100 U/ml penicillin (Gibco), and 2 mM glutamine (Gibco) at 37°C in 5 % CO_2_. Growth medium for BJ-5ta cells was further supplemented with hygromycin B (10 µg/ml, Invitrogen). PBMCs were obtained from healthy blood donors from the University Hospital Essen after informed consent under Ethical Review Board approvals 14-5961-BO and 16-6769-BO, in accordance with the Declaration of Helsinki. HLA typing was performed by next generation sequencing as described^72^. All blood donors were HCMV seropositive and were selected according their HLA-DPB1 and HLA-DRB1 typing matching the endogenously expressed HLA-DP and HLA-DR in HeLa cells.

### Viruses, infection, and HCMV lysate generation

Virus stocks from the HCMV strain AD169 ^64^, AD169-BAC2 ^73^, BAC2 ΔUS2-11 ^74^ and BAC20-EGFP ^75^ were generated as previously described by propagating HCMV in MRC-5 cells^76^. Viral titers were determined by standard plaque titration on MRC-5 cells. All infections were conducted with centrifugal enhancement (900 g for 30 min).

For generation of HCMV lysates for T-cell stimulation, BJ-5ta cells were infected with AD169-BAC2 at an MOI of 3 or mock treated. At 4 days post-infection, cells were scraped, washed and resuspended in PBS. After 5 freeze-thaw cycles, lysates were treated with ultra-sonication (2 times 10 sec) and centrifuged (1000 g, 20 min, 4°C). Supernatants were used for T-cell culture assays.

### Cytokine treatment

Human IFNγ (PBL), IFNα (PBL) and TNFα (PeproTech) were used in following concentrations: 200 U/ml, 200 U/ml and 20 ng/ml, respectively.

### Protein stability determination by cycloheximide chase assay

Protein stability was determined in transfected HeLa cells (see transfection, plasmids and mutagenesis) by treatment with 50 µg/ml cycloheximide (CHX, Roth). Cells were washed once in CHX-containing medium, followed by incubation in CHX-containing medium for indicated time periods. Finally, whole cell lysates were prepared and subjected to immunoblot analysis.

### Protein precipitation of cell culture supernatant

Cell culture supernatant of transiently transfected HeLa cells (see transfection, plasmids and mutagenesis) was collected. 400 µl supernatant were mixed with 1600 µl ice-cold acetone and incubated for 1 h at −20°C. After centrifugation for 90 min, 4°C and 13.000 g, the pellet was dried and resuspended in RIPA buffer. Subsequently, samples were subjected to immunoblot analysis.

### Transfection, plasmids, and mutagenesis

Transient transfection was performed using 1 or 2 µg plasmid DNA and 3.5 or 7 µl FuGENE HD transfection reagent (Promega) per 5x 10^5^ cells. Cells were transfected with the following plasmids: pUNO1-hCIITA (Invitrogen), pRP-humanCIITA-3xFLAG (VectorBuilder), pRP-3xFLAG-mouseCIITA (VectorBuilder), pcDNA3.1(+) (Invitrogen), pIRES_neo_-FLAG/HA-EYFP (RRID: Addgene_10825, Gift from Thomas Tuschl ^77^) and HCMV ORF Expression Library ^48^. Following plasmids were generated in this study: pcDNA3.1-US28HA, pcDNA3.1-US28HA-R129A, pcDNA3.1-US28HA-ΔC, pcDNA3.1-US27HA and pcDNA3.1-US29HA.

Cloning of pcDNA3.1-US28HA, pcDNA3.1-US27HA and pcDNA3.1-US29HA was performed using the primers listed in Supplementary Table S2. In order to generate US28 mutants, QuikChangeII XL Site-Directed Mutagenesis kit (Agilent Technologies) was used according to the manufacturer’s instructions with pcDNA3.1-US28HA plasmid as the template and respective primers (see Supplementary Table S2). All constructs were confirmed by DNA sequencing of the insert (LGC Genomics).

### Immunoblot analysis

For immunoblotting, whole cell lysates were prepared as described before in RIPA ^78^ or 5 M urea buffer and equal amounts of protein were subjected to SDS polyacrylamide gel electrophoresis (SDS-PAGE). Proteins were subsequently transferred onto nitrocellulose membranes and immunoblot analysis was performed with following antibodies: HA (Sigma-Aldrich, H6908), FLAG (M2, Sigma-Aldrich, F3165), β-tubulin (Cell Signaling, 2146), β-actin (Sigma-Aldrich, A2228), GAPDH (FL-335, Santa Cruz, sc-25778).. Proteins were visualized using peroxidase-coupled secondary antibodies (rabbit-POD, Sigma-Aldrich, A6154; mouse-POD, Jackson ImmunoResearch, 115-035-062) and an enhanced chemiluminescence system (Cell Signaling Technology).

### Immunoprecipitation

Cells were lysed (150 mM NaCl, 10 mM KCl, 10 mM MgCl_2_, 10 % [v/v] glycerol, 20 mM HEPES [pH 7.4], 0.5 % [v/v] NP-40, 0.1 mM phenylmethylsulfonyl fluoride [PMSF], 1 mM dithiothreitol [DTT], 10 µM pepstatin A, 5 µM leupeptin, 0.1 mM Na-vanadate, Complete protease inhibitor EDTA-free [Roche]). Lysates were centrifuged and immunoprecipitation (IP) antibody (anti-HA, HA-7, Sigma-Aldrich, H3663) was added to the supernatant. Precipitation of immune complexes was performed with protein G-Sepherose (GE Healthcare), benzonase (Sigma-Aldrich, E1014) digestion for 3 h at 4 °C, and washing steps with 150, 250 and 500 mM NaCl-containing buffer. Samples were further processed by immunoblot analysis.

### Semi-quantitative RT-PCR

For semi-quantitative RT-PCR, total RNA was isolated from 1x 10^6^ cells using the RNeasy Mini kit (Qiagen) and digested with DNase I. Subsequent one-step RT-PCR (Qiagen) was performed using gene-specific primers listed in Supplementary Table S2.

### Flow cytometry

For flow cytometry, cells were detached, washed with 2% FCS-PBS and stained with labelled antibodies. For intracytoplasmic staining, cells were fixed in 4% PFA-PBS for 15 min at room temperature after detachment and washing steps. Permeabilization was performed with 1% saponin-PBS for 15 min at room temperature, followed by antibody staining. The following antibodies were used: HLA-DP-BV421 (B7/21, BD Biosciences, 750875), HLA-DP-APC (B7/21, Leinco Technologies, H240), HLA-DP-PE (B7/21, BD Biosciences, 566825), HLA-DR-PE (L243, BD Biosciences, 347401), HLA-DQ-PE (HLADQ1, BioLegend, 318106), HLA-DM-PE (MaP.DM1, BD Biosciences, 555983), CD137-APC (4B4-1, BD Biosciences, 561702), CD4-PE-Cy7 (SK3, BD Biosciences, 557852), CD8-Pacific Blue (B9.11, Beckman Coulter, B49182), CD3-Krome Orange (UCHT1, Beckman Coulter, B00068), CD57-FITC (TB03, Miltenyi Biotec, 130-122-935). Measurements were performed in a Gallios 10/3 cytometer (Beckman Coulter), using the Kaluza for Gallios Acquisition software (Version 1.0, Beckman Coulter). Data analysis was conducted with FlowJow (Version 10.8.1, Tree Star).

### T-cell activation assay

To obtain HCMV-specific CD4+ T cells, PBMCs were pulsed with HCMV lysate (25 µg/ml) for 4 h at 37°C in 5% CO_2_. Cells were cultured for 14 days in RPMI 1640 (ccpro) supplemented with 10 % heat-inactivated human serum (Merck), 10 ng/ml IL-7, 1 ng/ml IL-12 and 5 ng/ml IL-15 (R&D Systems). Re-stimulation with irradiated (100 Gy), lysate-pulsed PBMCs was performed at day 7 of the co-culture. Afterwards, cells were cultured in RPMI 1640 (ccpro) supplemented with 10 % heat-inactivated human serum (Merck), 50 U/ml IL-2 (Miltenyi), 10 ng/ml IL-7 and 5 ng/ml IL-15 (R&D Systems). After 14 days, expanded T cells were re-challenged for 24 h with HeLa cells transfected with the respective plasmids and pulsed with mock or HCMV lysate (25 µg/ml). The specific T-cell response was quantified by flow cytometry as percentage of gated CD4+ T cells expressing the activation marker CD137 as previously described ^79^.

### Antiviral activity of T-cell supernatant

To evaluate the antiviral activity of supernatants from the T-cell activation assay, MRC-5 cells were incubated with these supernatants in serial dilutions for 24 h before cells were infected with HCMV-BAC20-EGFP at an MOI of 0.05. HCMV-induced EGFP expression was quantified using a Mithras^2^ LB 943 Multimode Reader (Berthold Technologies, Software MikroWin 2010). Microscopy was conducted with a Leica DM IL LED Microscope (Software LAS V4.0) and a Bioreader-7000 Fz (BIO-SYS, Software EazyReader).

### LS-MS/MS sample preparation

Cell lysates were processed according to the SP3 protocol ^80^ with minor modifications. Briefly, the cells were lysed in urea buffer (7M urea, 2M thiourea, 30 mM TRIS, 0.1 % sodium deoxycholate, pH 8.5) and an aliquot of 10 µg was reduced with dithiothreitol (5 mM final concentration, 50 °C, 15 min), and alkylated using 2-iodoacetamide (15 mM final concentration, RT, 15 min). Subsequently, 100 µg of SP3-beads were added and the volume was adjusted to 100 µL using 50 mM ammonium bicarbonate (ambic). 170 µL of acetonitrile (ACN) were added and samples were incubated for 18 min. After washing the beads twice with 180 µL 70 % EtOH and once with 180 µL ACN, 1 µg trypsin (SERVA Electrophoresis, Heidelberg, Germany) in 55 µL ambic was added and samples were digested overnight at 37 °C. Finally, the solution was transferred to a new vial, evaporated to dryness and peptides were resuspended in 100 µL 0.1 % trifluoracetic acid.

### Data-independent acquisition mass spectrometry

300 ng tryptic peptides per sample were analyzed in randomized order using a vanquish Neo UHPLC coupled to an Orbitrap 480 mass spectrometer (both Thermo Scientific). The mobile phase A consisted of 0.1 % formic acid (FA), mobile phase B of 80 % ACN and 0.1 % FA. Peptides were loaded on a trap column (Acclaim PepMap 100, 100 µm x 2cm, Thermo Scientific) using combined control with a loading volume of 20 µL, a maximum flow rate of 30 µL/min and a maximum pressure of 800 bar. Separation of peptides was achieved using a DNV PepMap Neo separation column (75 µm x 150 mm, Thermo Scientific) and a gradient from 1 % to 40 % B within 120 min and a flowrate of 400 nL/min at 60 °C. The MS parameters were set as follows: The RF lens amplitude was set to 55 %, the MS1 scan rage was 350-1450 m/z with a resolution of 120,000, a normalized AGC target of 300 % and a maximum injection time of 54 ms. MS2 scans were acquired using a resolution of 30,000, a HCD collision energy of 30 % and a scan rage of 145-1450 with a normalized AGC target of 2,500 % and a maximum injection time of 80 ms. A total of 40 isolation windows between 350 and 1450 m/z were cycled through with one MS1 scan being recorded after every 21 MS2 scans.

### Mass spectrometry data analysis

Protein identification and quantification was conducted using DIA-NN (v.1.8.1; ^81^) in library-free mode. The SwissProt database restricted to Homo sapiens as well as the Uniprot reference proteome for the Human cytomegalovirus (both ver. 2022_02) were used for peptide identification. Default settings were used, except for the neural network classifier, which was used in double-pass mode, and protein inference, which was set to species-specific. The report file was filtered for all q-values ≤ 0.01 using R (ver. 4.3.0; www.r-project.org). Subsequently, protein quantities were calculated using the MaxLFQ algorithm as implemented in the DIA-NN R package. Missing data was imputed on protein level using the mixed imputation function from the imp4p package ^82^. Statistically significant differences between the experimental groups were assessed by means of ANOVA followed by Tukey’s HSD posthoc tests. The ANOVA p-value was corrected for multiple testing according to the method of Benjamini-Hochberg. The significance threshold was set to pFDR ≤ 0.05, p posthoc ≤ 0.05 and a ratio of mean intensities ≥ 2 or ≤ 0.5.

### Quantification and statistical analysis

The resulting data were analyzed using GraphPad Prism software. The values are reported as Mean ± standard deviation (SD). Statistical significance was tested by applying the respective test indicated in the figure legends.

---

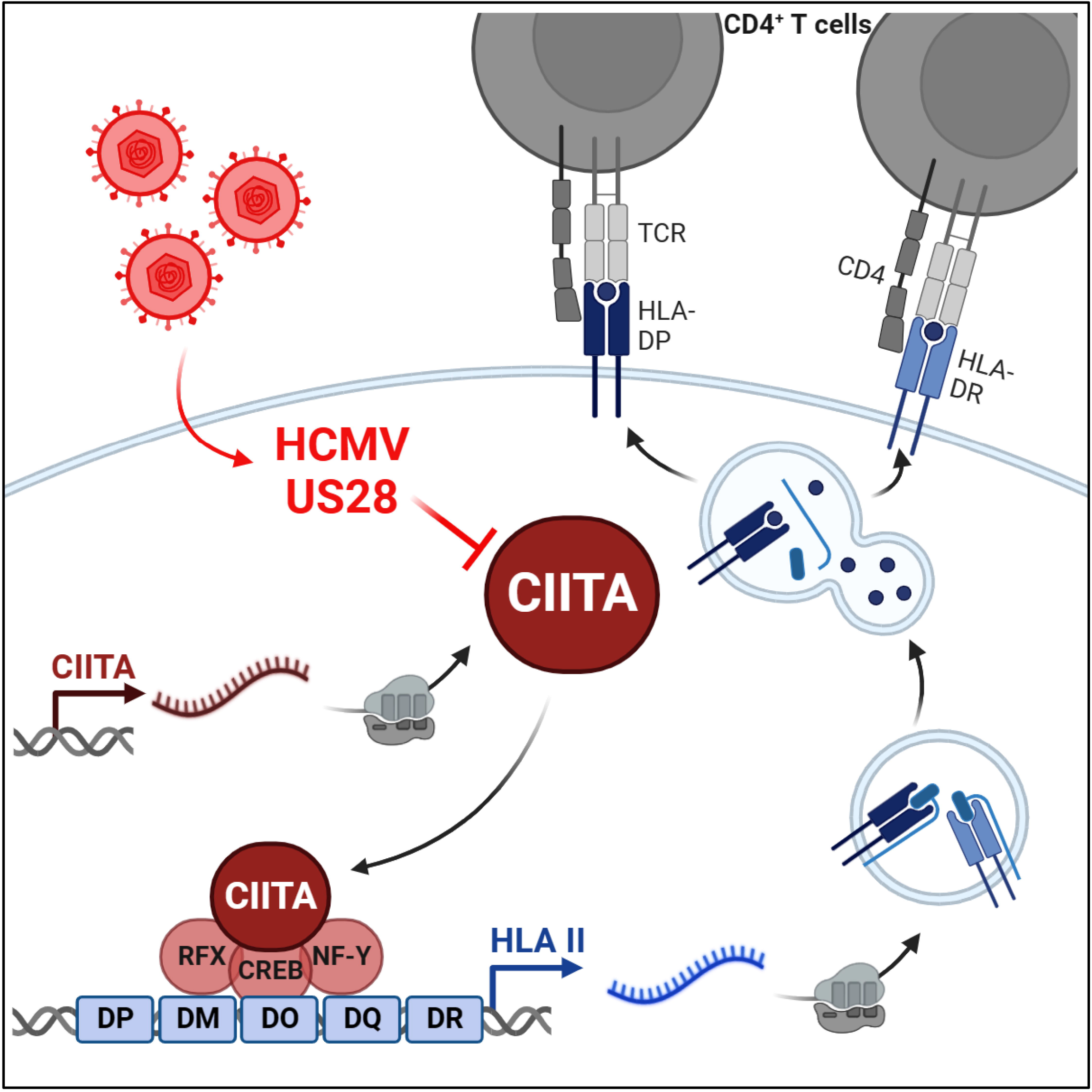

