## Supplementary material for "The human cytomegalovirus-encoded pUS28 antagonizes CD4+ T-cell recognition by targeting CIITA": Suppl.-Table S2 Primers

Supplementary Table S2: Primer sequences

| | Forward | Reverse |
|---|---|---|
| Cloning HCMV-US28HA | CGGCTAGCATGACACCGACGACGACGACC | CGCTCGAGTTAAGCGTAATCTGGAACATCGTATGGGTACGGTATAATTTGTGAGACGCG |
| Cloning HCMV-US27HA | CGGCTAGCATGACCACCTCTACAAATAATC | CGCTCGAGTTAAGCGTAATCTGGAACATCGTATGGGTACAACAGAAATTCCTCCTCCCC |
| Cloning HCMV-US29HA | CGGCTAGCATGCGGTGTTTCCGATGGTGG | CGGAATTCTTAAGCGTAATCTGGAACATCGTATGGGTACTCGGAGGTGTCAACAACCC |
| QuikChange US28-R129A | CACGGAGATTGCACTCGATGCCTACTACGCTATTGTTTAC | GTAAACAATAGCGTAGTAGGCATCGAGTGCAATCTCCGTG |
| QuikChange US28-ΔC (F298-Stop) | CTTCGTGGGCACCAAGTACCCATACGATGTTCCAGATTACGCTTAGCGGCAAGAACTACACTG | CAGTGTAGTTCTTGCCGCTAAGCGTAATCTGGAACATCGTATGGGTACTTGGTGCCCACGAAG |
| HLA-DPB1 (Meurer et al., 2018) | GCTTCCTGGAGAGATACATC | CAGCTCGTAGTTGTGTCTGC |
| HLA-DR (Morimoto et al., 2004) | GCCAACCTGGAAATCATGAC | AGGGCTGTTCGTGAGCACA |
| CIITA (Sandhu et al., 2020) | AGCCTTTCAAAGCCAAGTCC | TTGTTCTCACTCAGCGCATC |
