## Supplementary material for "The human cytomegalovirus-encoded pUS28 antagonizes CD4+ T-cell recognition by targeting CIITA": Suppl.-Fig. S1-S4

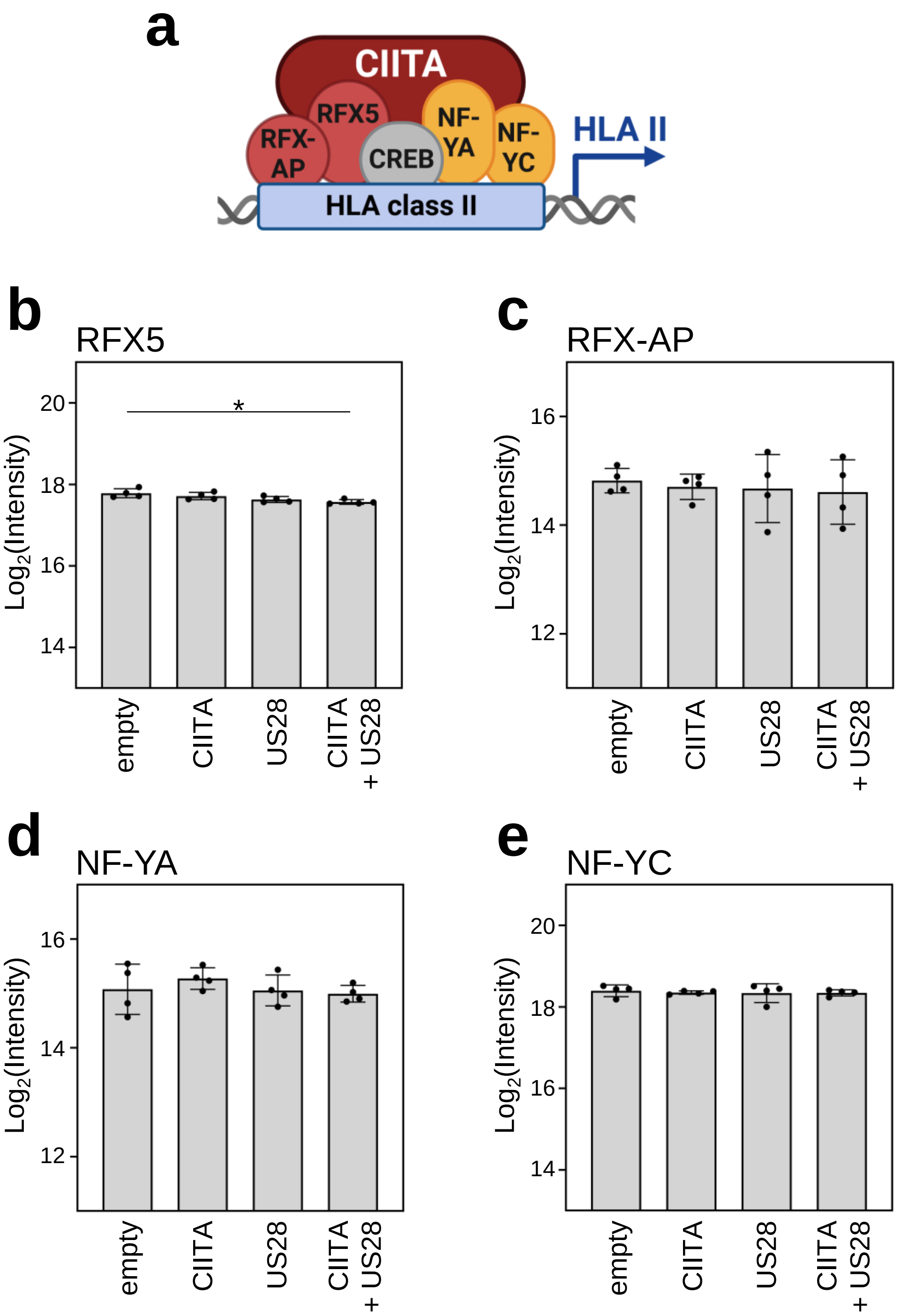


Supplementary Figure S1:

**Global proteome analysis showed that the other components of the CIITA enhanceosome are constitutively expressed**

(a) shows a simplified schema of the CIITA enhanceosome. (b) to (e) depict changes in the abundance of selected proteins measured by MS: (b) RFX5, (c) RFX-AP, (d) NF-YA, (e) NF-YC as log2 (intensity) values of untreated cells, cells expressing either CIITA, pUS28 or both (n = 4).

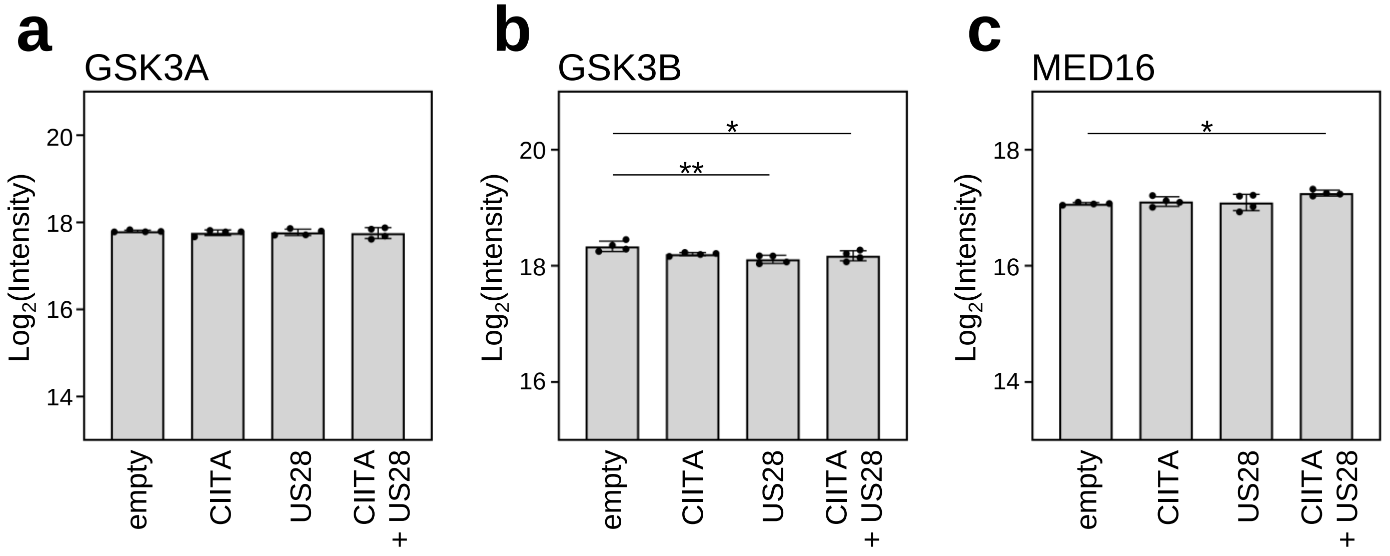


Supplementary Figure S2:

**Global proteome analysis showed that recently described regulators of HLA-II transcription are constitutively expressed**

Changes in the abundance of selected proteins measured by MS: (a) GSK3A, (b) GSK3B, (c) MED16 as log2 (intensity) values of untreated cells, cells expressing either CIITA, pUS28 or both (n = 4).

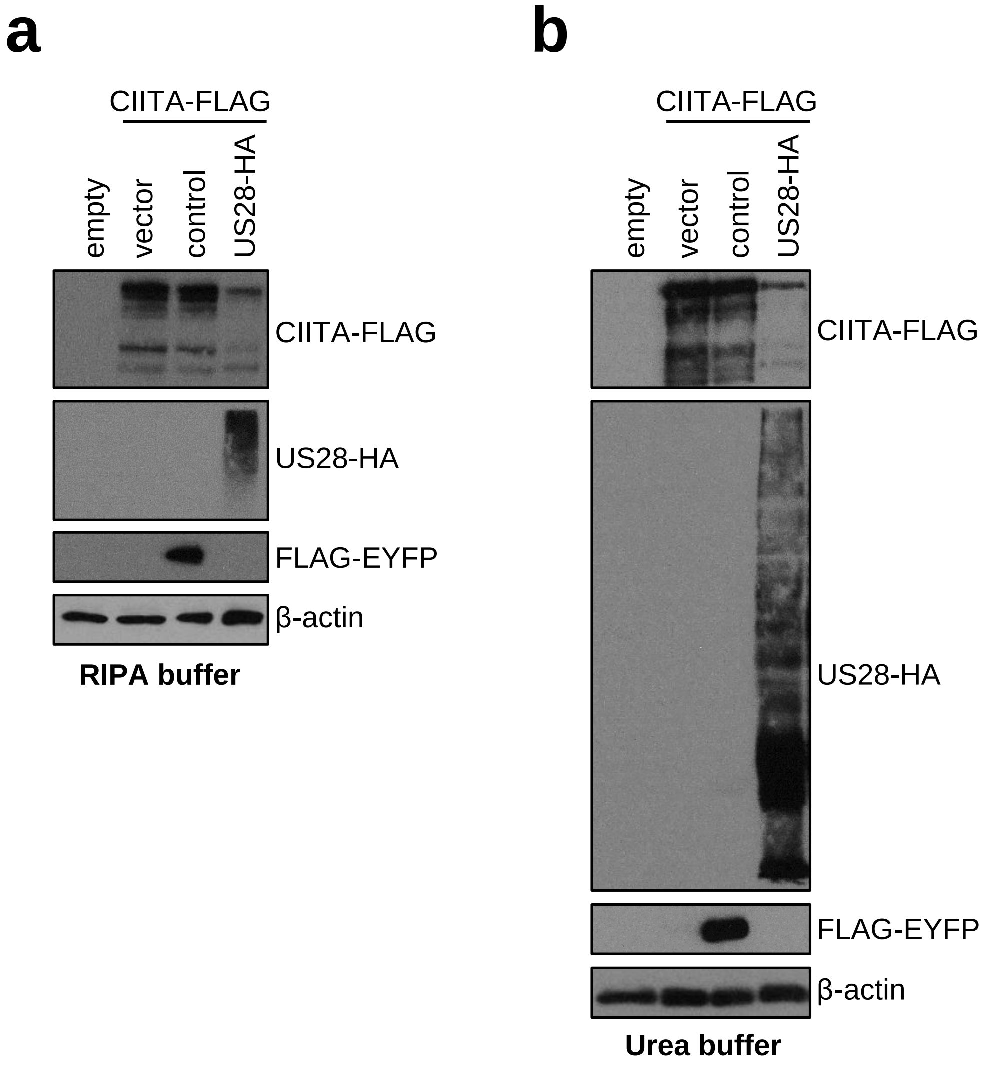


Supplementary Figure S3:

**pUS28 diminishes CIITA protein levels also in denaturing lysis buffer**

HeLa cells were either left untreated or were co-transfected with CIITA-3xFLAG expression construct and pcDNA:US28-HA, pIRESNeo-FLAG/HA-EYFP (control) or empty vector. At 24 h post-transfection, protein lysates were prepared in RIPA (a) or denaturing lysis buffer based on 5 M urea (b) and analyzed by immunoblot using antibodies detecting the indicated proteins or the respective epitope tags.

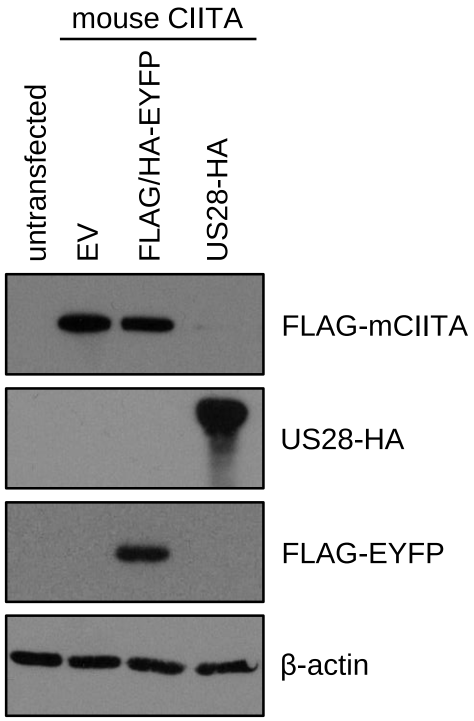


Supplementary Figure S4:

**pUS28 diminishes protein levels of murine CIITA**

HeLa cells were either left untreated or were co-transfected with 3xFLAG-mouseCIITA expression construct and pcDNA:US28-HA, pIRESNeo-FLAG/HA-EYFP (control) or empty vector. At 24 h post-transfection, protein lysates were prepared and analyzed by immunoblot using antibodies detecting the indicated proteins or the respective epitope tags.
